# G_2_C_4_ targeting antisense oligonucleotides potently mitigate TDP-43 dysfunction in C9orf72 ALS/FTD human neurons

**DOI:** 10.1101/2023.06.26.546581

**Authors:** Jeffrey D. Rothstein, Victoria Baskerville, Sampath Rapuri, Emma Mehlhop, Paymaan Jafar-nejad, Frank Rigo, Frank Bennett, Sarah Mizielinska, Adrian Isaacs, Alyssa N. Coyne

## Abstract

The G_4_C_2_ repeat expansion in the C9orf72 gene is the most common genetic cause of Amyotrophic Lateral Sclerosis and Frontotemporal Dementia. Many studies suggest that dipeptide repeat proteins produced from this repeat are toxic, yet, the contribution of repeat RNA toxicity is under investigated and even less is known regarding the pathogenicity of antisense repeat RNA. Recently, two clinical trials targeting G_4_C_2_ (sense) repeat RNA via antisense oligonucleotide failed despite a robust decrease in sense encoded dipeptide repeat proteins demonstrating target engagement. Here, in this brief report, we show that G_2_C_4_ antisense, but not G_4_C_2_ sense, repeat RNA is sufficient to induce TDP-43 dysfunction in induced pluripotent stem cell (iPSC) derived neurons (iPSNs). Unexpectedly, only G_2_C_4_, but not G_4_C_2_ sense strand targeting, ASOs mitigate deficits in TDP-43 function in authentic C9orf72 ALS/FTD patient iPSNs. Collectively, our data suggest that the G_2_C_4_ antisense repeat RNA may be an important therapeutic target and provide insights into a possible explanation for the recent G_4_C_2_ ASO clinical trial failure.

## Introduction

An intronic GGGGCC (G_4_C_2_) hexanucleotide repeat expansion (HRE) in the C9orf72 gene is the most common cause of familial ALS and FTD, collectively referred to as C9orf72 ALS/FTD, and also causes ∼8% of apparently sporadic ALS or FTD ^1,2^. The HRE is bidirectionally transcribed to form G_4_C_2_ (sense) and G_2_C_4_ (antisense) RNA species both of which have been documented to pathologically accumulate into RNA foci in human autopsy tissue and induced pluripotent stem cell (iPSC) lines, as well as in mouse and fly models of C9orf72 ALS/FTD. In addition, repeat-associated non-ATG (RAN) translation of G_4_C_2_ and G_2_C_4_ RNA produces five distinct dipeptide repeat (DPR) proteins: sense encoded Poly(GA), Poly(GP), Poly(GR) and antisense encoded Poly(PR), Poly(PA), Poly(GP). Together, these RNA species and DPR proteins are thought to contribute to disease through gain of toxicity mechanisms ^3^. While much research has concluded that specific DPRs, including Poly(PR) and Poly(GR), translated from the G_2_C_4_ and G_4_C_2_ repeat RNA strands respectively, are toxic when highly overexpressed in multiple model systems ^4-8^, little is known regarding the pathogenicity of repeat RNAs themselves especially in endogenous expression models where it has been technically challenging to dissociate repeat RNA and DPR mediated toxicity.

It is thought that expression of C9orf72 repeat RNA species can sequester RNA binding proteins (RBPs) into pathologic RNA foci and impede their function ^9,10^ and disrupt nuclear pore complexes and nucleocytoplasmic transport ^10,11^. Importantly, like 97% of ALS and ∼50% of FTD cases, the nuclear clearance along with associated loss of nuclear function and subsequent cytoplasmic mislocalization of the RNA binding protein TDP-43 is a common pathological hallmark of C9orf72 ALS/FTD ^12,13^. However, early histology studies failed to evaluate the functional consequences of this interaction as well as the role of soluble repeat RNA species in disease pathogenesis. In fact, the vast number of studies published in the last decade have focused solely on the DPR species. Notably, prior work from ourselves and others have shown that the G_4_C_2_ sense repeat RNA can be itself toxic in human neurons ^11^. However, little is known regarding the pathobiology of the G_2_C_4_ antisense RNA species in human neurons. Although antisense pathology is detectable in CNS regions at autopsy, multiple reports suggest that the expression of G_2_C_4_ repeat RNA and presence of G_2_C_4_ foci is low in human tissue, as compared to G_4_C_2_ RNA. However, the exact length and proportion of these different transcripts remains unclear. Furthermore, histological pathology, by itself, is not a measure of physiological toxicity. Nonetheless, the reliable presence of translated DPR products from the antisense strand (e.g. Poly(PR) and Poly(PA)) provides clear *in vivo* evidence of protein products from antisense RNA in human brain and spinal cord ^1,2,14-20^. Interestingly, histological analyses in postmortem C9orf72 ALS/FTD patient tissues have suggested that nuclear depletion and the mislocalization of the RNA binding protein TDP-43 from the nucleus to the cytoplasm is more frequently observed in cells containing abundant G_2_C_4_ antisense repeat RNA foci ^14,21^. Thus, these studies highlight relationship between G_2_C_4_ antisense repeat RNA and TDP-43 pathology in C9orf72 disease which may be therapeutically relevant and worthy of functional studies.

Due to the vast literature largely focused on sense strand RNA abundance and sense strand DPR products, antisense oligonucleotide (ASO) therapies targeting the G_4_C_2_ sense strand repeat RNA have been identified. Unfortunately, an international clinical trial of the G_4_C_2_ (sense) targeting ASO BIIB078 (NCT04288856) in over 100 C9orf72 ALS patients was terminated (https://investors.biogen.com/news-releases/news-release-details/biogen-and-ionis-announce-topline-phase-1-study-results) due to lack of efficacy despite a robust BIIB078 initiated reduction in G_4_C_2_ sense encoded DPRs in patient CSF. Similarly, Wave Pharmaceuticals terminated their C9orf72 ASO trial, which also targeted the sense strand, due to lack of efficacy (https://www.thepharmaletter.com/article/wave-life-sciences-ends-wve-004-program). Importantly, these G_4_C_2_ sense strand targeting ASO therapies have no known effect on the G_2_C_4_ antisense RNA or its DPR products. While ASOs targeting G_4_C_2_ sense repeat RNA can partially reduce G_4_C_2_ linked DPR pathology ^11,22-24^, the therapeutic potential of targeting the G_2_C_4_ antisense repeat RNA is completely unknown in human systems. As such, an understanding of the G_2_C_4_ repeat RNA elicited toxicity is critically needed.

In this brief report, we utilize iPSNs to demonstrate that G_2_C_4_ but not G_4_C_2_ repeat RNA expression is sufficient to induce a loss of nuclear TDP-43 function in human spinal neurons. Moreover, using antisense oligonucleotides that specifically target G_4_C_2_ or G_2_C_4_ repeat RNAs, we find that reduction of antisense but not sense repeat RNA restores TDP-43 function in authentic C9orf72 ALS/FTD patient iPSNs expressing endogenous levels of the C9orf72 repeat expansion. Together, our data suggest that therapeutic targeting of G_2_C_4_ antisense repeat RNA may be beneficial for restoring TDP-43 function in C9orf72 ALS/FTD.

## Results

Nuclear clearance and cytoplasmic mislocalization and aggregation of the RNA binding protein is a pathological hallmark of ALS and related neurodegenerative diseases ^12,13^. It is thought that nuclear loss of TDP-43 function is an early and significant contributor to neurodegenerative disease pathogenesis ^25-29^ and histology studies have suggested that nuclear loss of TDP-43 precedes cytoplasmic aggregation ^13^. Consistent with the hypothesis that nuclear loss of TDP-43 function precedes cytoplasmic aggregation, studies in C9orf72 iPSNs have demonstrated the emergence of a molecular signature of TDP-43 dysfunction prior to overt nuclear clearance and cytoplasmic mislocalization ^11,30^. Interestingly, prior analyses have reported that TDP-43 pathology is more prevalent in CNS cells with G_2_C_4_ repeat RNA foci compared to those with G_4_C_2_ foci ^14,21^. Thus, we hypothesized that G_2_C_4_ repeat RNA itself may contribute to TDP-43 loss of function observed in C9orf72 ALS/FTD. To address the contribution of G_2_C_4_ repeat RNA to TDP-43 dysfunction and mislocalization in human neurons, we first utilized repeat RNA-only constructs previously demonstrated to produce G_4_C_2_ and G_2_C_4_ repeat RNA but not DPRs ^31,32^ and generated an additional long G_2_C_4_ repeat construct capable of expression in mammalian cells. Recent studies have identified a number of reliable gene expression and RNA processing changes that occur following TDP-43 depletion in human neurons ^25,26,28^. We have recently demonstrated that a panel of these RNA targets and associated splicing alterations can be employed as a robust measure of TDP-43 functionality in C9orf72 ALS/FTD and sALS iPSNs ^30^. Using qRT-PCR to evaluate the expression of multiple TDP-43 mRNA targets, we observe a significant molecular hallmark of TDP-43 dysfunction, including the emergence of multiple cryptic exon containing mRNA species, following the expression of G_4_C_2_ and G_2_C_4_ repeat RNA-only constructs in otherwise wildtype iPSNs (**Figure 1**). Consistent with a loss of nuclear TDP-43 function, immunostaining and confocal imaging revealed a small but statistically significant number of G_2_C_4_, but not G_4_C_2_ expressing neurons that contained observable new cytoplasmic TDP-43 immunoreactivity (**Supplemental Figure 1**). However, we note that this cytoplasmic signal was minimal suggesting that TDP-43 nuclear function may be impacted prior to overt histological mislocalization. Collectively, these data indicate that G_2_C_4,_ repeat RNA expression is sufficient to induce TDP-43 dysfunction in human neurons and suggest that perhaps our prior observations of molecular hallmarks of TDP-43 loss of function in endogenous C9orf72 iPSNs ^30^ could be predominantly linked to endogenous antisense G_2_C_4_ and not G_4_C_2_ repeat RNA expression.

**Figure 1:**
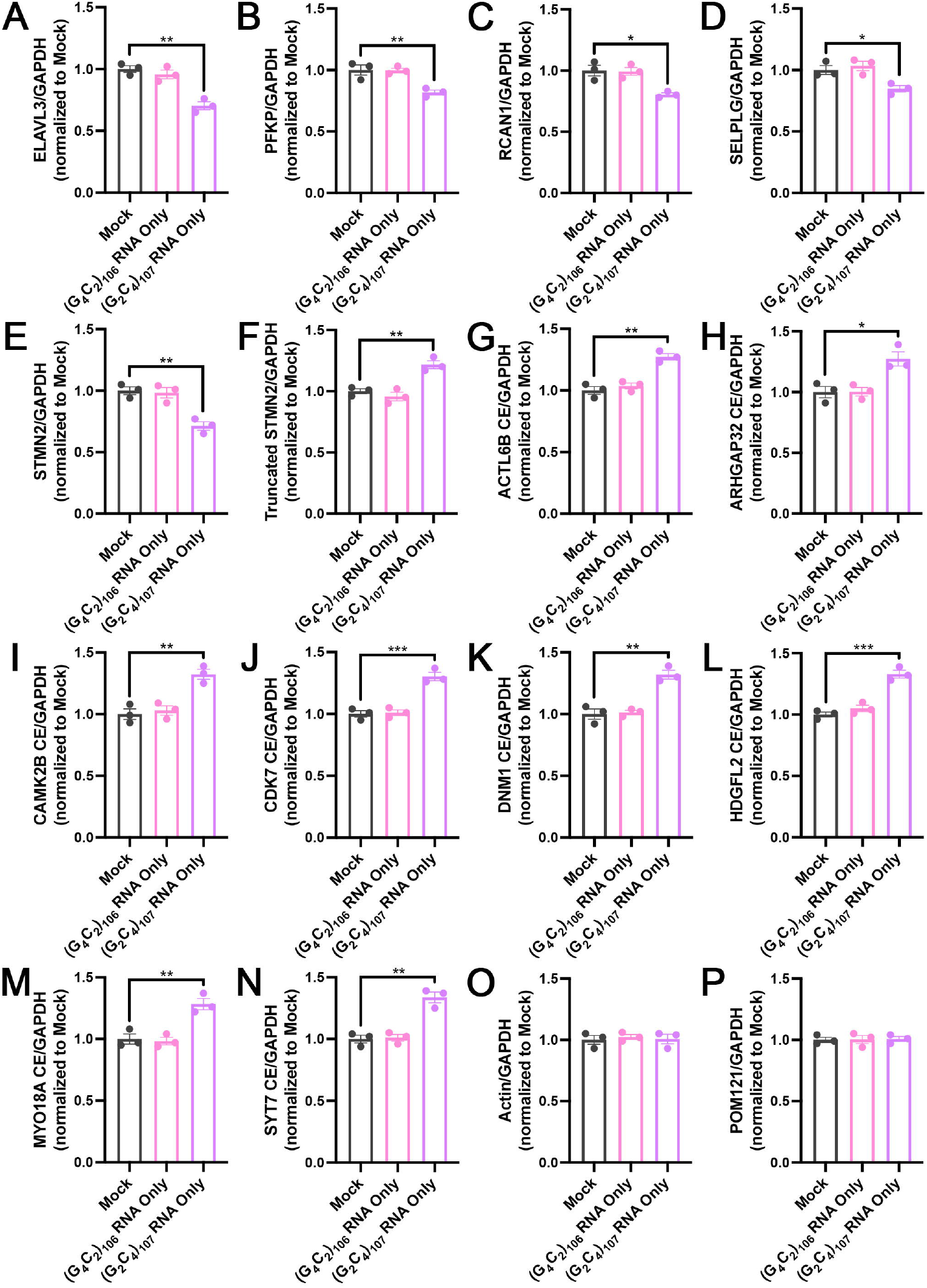
G_2_C_4_ antisense repeat RNA expression triggers TDP-43 dysfunction in iPSNs. (**A-P**) qRT-PCR for *ELAVL3* (**A**), *PFKP* (**B**), *RCAN1* (**C**), and *SELPLG* (**D**), *STMN2* (**E**), truncated *STMN2* (**F**), *ACTL6B* cryptic exon containing (**G**), *ARHGAP32* cryptic exon containing (**H**), *CAMK2B* cryptic exon containing (**I**), *CDK7* cryptic exon containing (**J**), *DNM1* cryptic exon containing (**K**), *HDGFL2* cryptic exon containing (**L**), *MYO18A* cryptic exon containing (**M**), *SYT7* cryptic exon containing (**N**), *POM121* (**O**), and *ACTIN* (**P**) mRNA in control iPSNs 2 weeks following expression of G_4_C_2_ or G_2_C_4_ repeat RNA only plasmids. GAPDH was used for normalization. *POM121* and *ACTIN* were used as negative control mRNAs not known to be regulated by TDP-43. n = 3 control iPSC lines. One-way ANOVA with Tukey’s multiple comparison test was used to calculate statistical significance. * p < 0.05, ** p < 0.01, *** p < 0.001.

In C9orf72 neurodegeneration, ASOs have previously been employed to induce the highly selective degradation of *C9orf72* mRNA or G_4_C_2_ repeat RNA ^22^. To facilitate the study of sense vs antisense repeat RNA toxicity in iPSNs expressing endogenous levels of the C9orf72 HRE, ASOs were designed to selectively target G_4_C_2_ (S-ASO) or G_2_C_4_ (AS-ASO) repeat containing RNAs. To test the efficacy of these ASOs in the selective degradation of G_4_C_2_ or G_2_C_4_ repeat RNA, we performed qRT-PCR for G_4_C_2_ or G_2_C_4_ repeat RNA in C9orf72 iPSC derived spinal neurons following 5, 10, 15, and 20 days of treatment with ASOs. We observed a significant strand specific decrease in G_4_C_2_ and G_2_C_4_ repeat RNA expression as early as 5 days after treatment (**Supplemental Figure 2**), an observation that is supported by prior reports that G_4_C_2_ repeat RNA knockdown happens significantly faster than DPR reduction ^11,22,24^. Consistent with this notion, we recently established that 5 day treatment with ASOs targeting G_4_C_2_ sense repeat RNA has no impact on Poly(GP) levels ^11^. Important for evaluation of biological specificity, the G_4_C_2_ targeting ASO had no impact on G_2_C_4_ expression and the ASO targeting G_2_C_4_ containing RNA had no impact on G_4_C_2_ expression (**Supplemental Figure 2**).

Having observed specificity for G_4_C_2_ and G_2_C_4_ repeat RNA targeting ASOs, we next asked whether repeat RNA strand specific ASO treatment could alleviate disease pathophysiology, namely TDP-43 dysfunction, in human iPSNs derived from multiple C9orf72 patients. Although we have previously reported that G_4_C_2_ repeat RNA ASO treatment can *prevent* cytotoxicity in C9orf72 iPSNs ^11,22^, importantly for this study we elected to initiate ASO treatment *after* the emergence of TDP-43 dysfunction, reminiscent of clinical treatment paradigms. To determine whether G_4_C_2_ and G_2_C_4_ targeting ASO treatment could restore TDP-43 function, we performed qRT-PCR for a panel of TDP-43 mRNA targets in C9orf72 iPSNs treated with strand specific ASOs. In doing so, we found that 15 day treatment with G_2_C_4_ antisense, *but not* G_4_C_2_ sense, repeat RNA targeting ASOs was sufficient to restore the expression of TDP-43 mRNA targets (**Figure 2**) thereby alleviating TDP-43 dysfunction. Consistent with a restoration of TDP-43 function, we observed a significant reduction in the percentage of C9orf72 iPSNs with observable cytoplasmic TDP-43 immunoreactivity following 20 days of treatment with G_2_C_4_ but not G_4_C_2_ repeat RNA targeting ASOs (**Supplemental Figure 3**). Interestingly, immunoprecipitation and qRT-PCR experiments indicate that G_2_C_4_ antisense but not G_4_C_2_ sense repeat RNA is enriched in TDP-43 protein-RNA complexes in C9orf72 patient iPSNs (**Supplemental Figure 4**) further underscoring the relationship between G_2_C_4_ antisense repeat RNA and TDP-43 in C9orf72 ALS/FTD pathogenesis.

**Figure 2:**
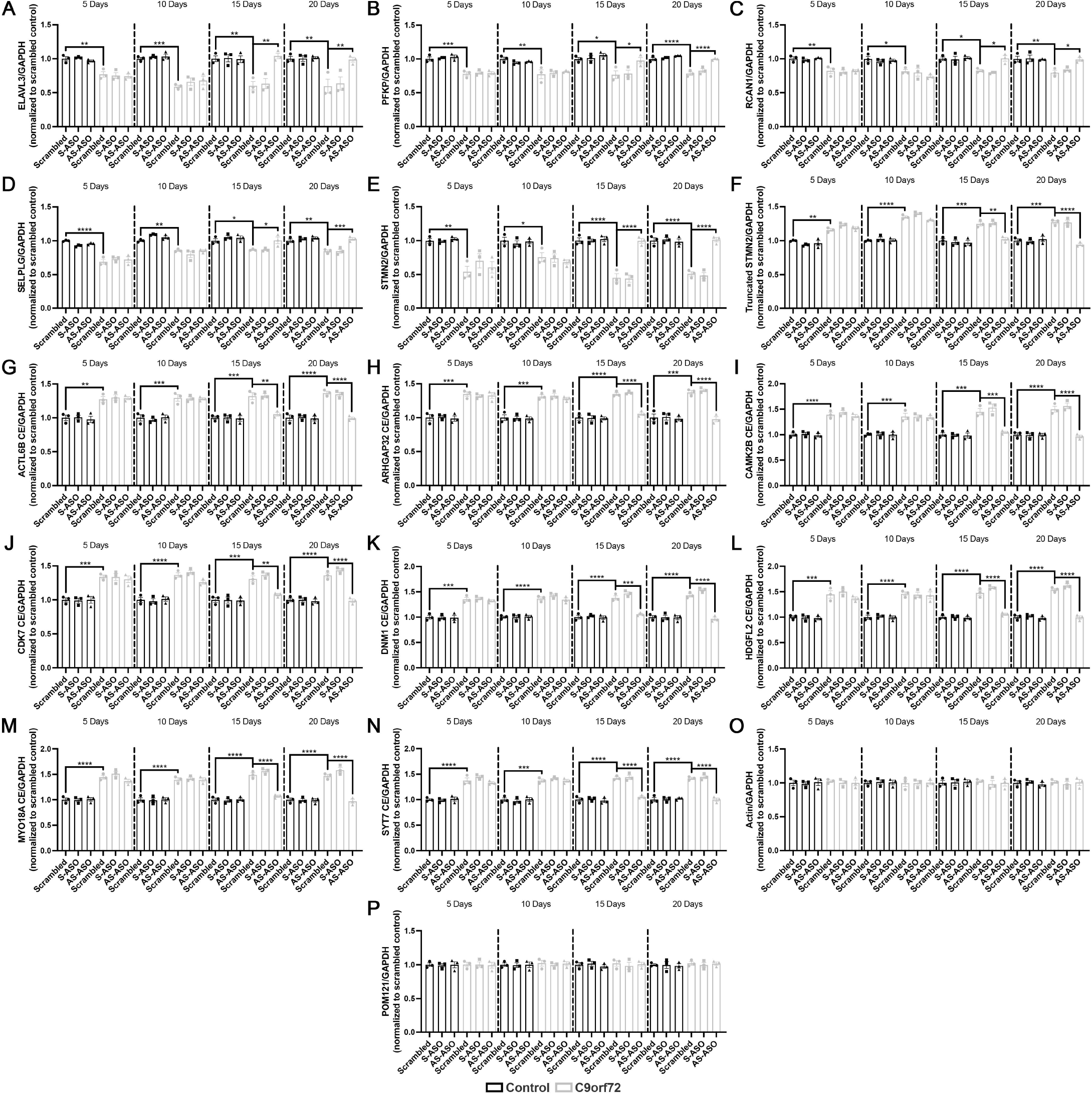
G_2_C_4_ targeting ASO treatment alleviates TDP-43 dysfunction in C9orf72 patient iPSNs. (**A-P**) qRT-PCR for *ELAVL3* (**A**), *PFKP* (**B**), *RCAN1* (**C**), and *SELPLG* (**D**), *STMN2* (**E**), truncated *STMN2* (**F**), *ACTL6B* cryptic exon containing (**G**), *ARHGAP32* cryptic exon containing (**H**), *CAMK2B* cryptic exon containing (**I**), *CDK7* cryptic exon containing (**J**), *DNM1* cryptic exon containing (**K**), *HDGFL2* cryptic exon containing (**L**), *MYO18A* cryptic exon containing (**M**), *SYT7* cryptic exon containing (**N**), *POM121* (**O**), and *ACTIN* (**P**) mRNA in control and C9orf72 iPSNs 5, 10, 15, and 20 days following a single dose of scrambled or repeat targeting ASO. GAPDH was used for normalization. *POM121* and *ACTIN* were used as negative control mRNAs not known to be regulated by TDP-43. n = 3 control and 3 C9orf72 iPSC lines. Two-way ANOVA with Tukey’s multiple comparison test was used to calculate statistical significance. * p < 0.05, ** p < 0.01, *** p < 0.001, **** p < 0.0001.

We next tested the hypothesis that combined treatment with G_4_C_2_ and G_2_C_4_ repeat RNA targeting ASOs might enhance the therapeutic benefit of G_2_C_4_ repeat RNA targeting ASO treatment alone, at least in regard to TDP-43 functionality. Importantly, the efficacy of repeat RNA knockdown was essentially unchanged following simultaneous treatment with both repeat RNA targeting ASOs (**Supplemental Figure 5**). Importantly, using qRT-PCR, we did not observe an increased benefit of TDP-43 function restoration upon simultaneous treatment with G_4_C_2_ and G_2_C_4_ targeting ASOs compared to G_2_C_4_ targeting ASOs alone (**Figure 3**). These data suggest that there is no added benefit of targeting both sense and antisense C9orf72 repeat RNAs to alleviate TDP-43 dysfunction in human neurons.

**Figure 3:**
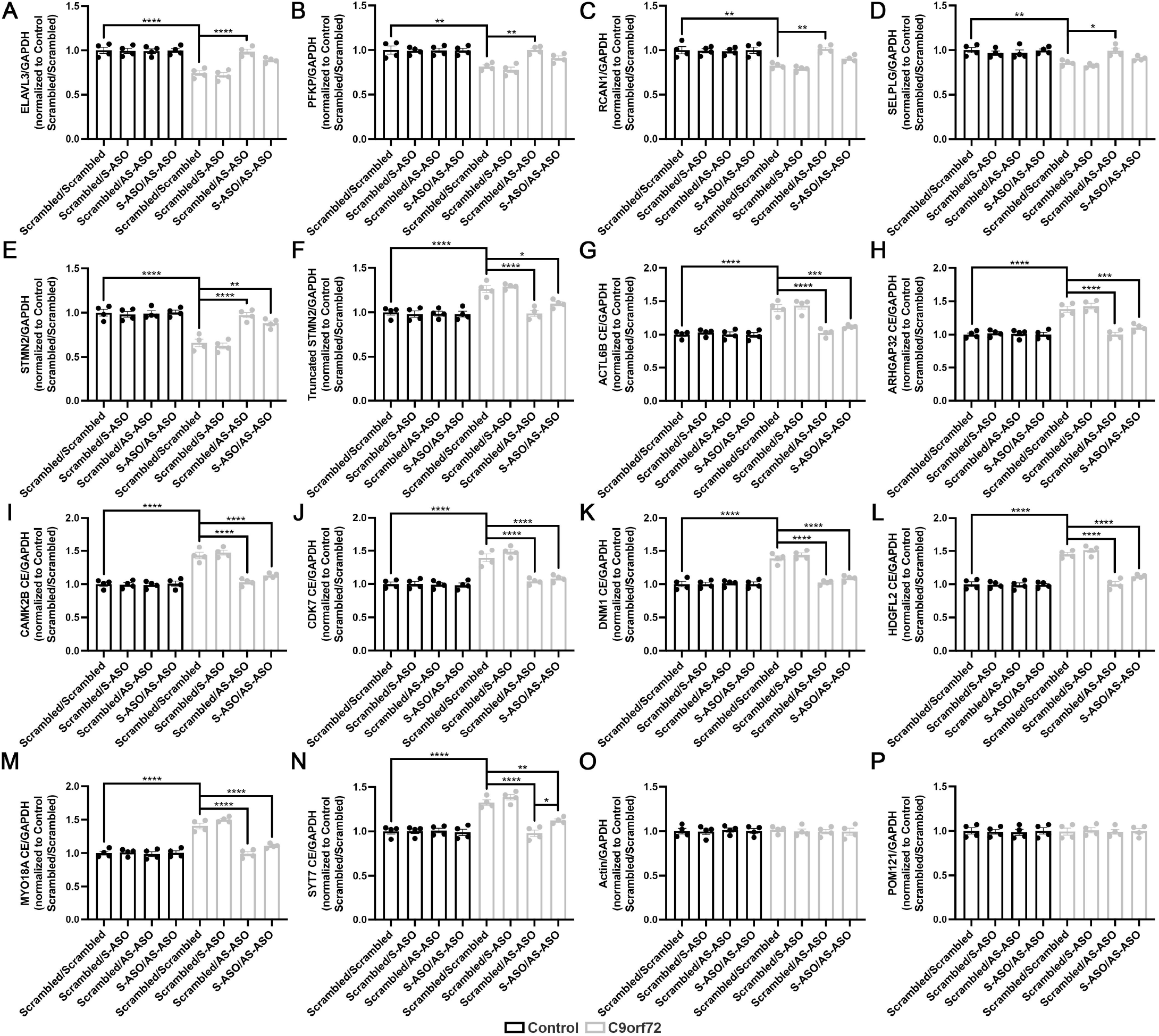
Simultaneous treatment with G_4_C_2_ and G_2_C_4_ repeat RNA targeting ASOs does not enhance the protective effects of G_2_C_4_ targeting ASOs alone. (**A-P**) qRT-PCR for *ELAVL3* (**A**), *PFKP* (**B**), *RCAN1* (**C**), and *SELPLG* (**D**), *STMN2* (**E**), truncated *STMN2* (**F**), *ACTL6B* cryptic exon containing (**G**), *ARHGAP32* cryptic exon containing (**H**), *CAMK2B* cryptic exon containing (**I**), *CDK7* cryptic exon containing (**J**), *DNM1* cryptic exon containing (**K**), *HDGFL2* cryptic exon containing (**L**), *MYO18A* cryptic exon containing (**M**), *SYT7* cryptic exon containing (**N**), *POM121* (**O**), and *ACTIN* (**P**) mRNA in control and C9orf72 iPSNs 15 days following 5 μM dosing with a combination of scrambled, G_4_C_2_, and G_2_C_4_ repeat RNA targeting ASOs. GAPDH was used for normalization. *POM121* and *ACTIN* were used as negative control mRNAs not known to be regulated by TDP-43. n = 4 control and 4 C9orf72 iPSC lines. Two-way ANOVA with Tukey’s multiple comparison test was used to calculate statistical significance. * p < 0.05, ** p < 0.01, *** p < 0.001, **** p < 0.0001.

## Discussion

Reduced nuclear localization of TDP-43 and corresponding loss of nuclear TDP-43 function is widely regarded as a prominent pathophysiological event contributing to neuronal demise in ALS, FTD, and related neurodegenerative diseases ^25-29^. To date, therapeutic strategies are aimed at restoring the expression of singular TDP-43 mRNA targets such as STMN2 ^26,27^ as opposed to broadly alleviating TDP-43 dysfunction. Further, in the case of C9orf72 ALS/FTD, although G_2_C_4_ antisense repeat RNA foci correlate with TDP-43 pathology at end stage disease ^14,21^, little is known regarding the contribution of pathologic C9orf72 repeat RNA species to TDP-43 dysfunction in authentic human neurons. In this brief report, we have demonstrated that G_2_C_4_ antisense, but not G_4_C_2_ sense, repeat RNA compromises TDP-43 functionality in endogenous C9orf72 human neurons. Although the mechanism by which G_2_C_4_ antisense repeat RNA specifically elicits TDP-43 dysfunction remains unknown and will require further future studies, this pathophysiological consequence is consistent with postmortem histological analyses indicating a correlation between G_2_C_4_ repeat RNA foci and TDP-43 pathology ^14,21^. Our RNA immunoprecipitation experiments (**Supplemental Figure 4**) suggest that TDP-43 may be at least partially sequestered away from it’s multiple RNA targets by G_2_C_4_ repeat RNA. Further, our data highlight the possibility that G_2_C_4_ repeat RNA induced TDP-43 dysfunction occurs in the absence of and perhaps prior to overt TDP-43 nuclear clearance and mislocalization to the cytoplasm (**Figure 1, Supplemental Figure 1**). We note that we and others have been unable to detect the most well-characterized mouse TDP-43 associated RNA misprocessing event, altered sortilin splicing ^33^, in C9orf72 BAC ^34^ or AAV9-(G_4_C_2_)_149_ transgenic mice ^35^ (unpublished data). These mouse models also do not show histological evidence of actual nuclear TDP-43 clearing typical of sporadic ALS or C9orf72 ALS/FTD (^34,35^, unpublished data). Thus, at the current time, this highlights authentic C9orf72 patient iPSNs as an essential model for evaluating the efficacy of therapeutic strategies in the repair of TDP-43 functionality in ALS/FTD.

Importantly, our data illuminate new insights into repeat RNA pathophysiology in C9orf72 ALS/FTD and provide human relevant data suggesting that G_2_C_4_ repeat RNA could be a significant contributor to disease pathogenesis. Collectively, these data suggest that targeting G_2_C_4_ antisense repeat RNA may be a viable therapeutic strategy for the restoration of TDP-43 function in ALS/FTD. Further, our data may provide insights into one reason that G_4_C_2_ sense stand targeting C9orf72 therapy failed to have clinical efficacy. However, these studies in authentic human C9orf72 ALS/FTD iPSNs, do not support the possibility of a combined sense and antisense repeat RNA targeting ASO therapy at least for the alleviation of TDP-43 dysfunction.

## Materials and Methods

### iPSC Derived Neuronal Differentiation

C9orf72 and non-neurological control iPSC lines were obtained from the Answer ALS repository at Cedars-Sinai (see **Supplemental Table 1** for demographics), maintained according to Cedars Sinai SOP, and differentiated into spinal neurons as previously described ^11,30,36^. All cells were maintained at 37°C with 5% CO_2_. iPSCs and iPSNs routinely tested negative for mycoplasma.

**Supplemental Table 1:**
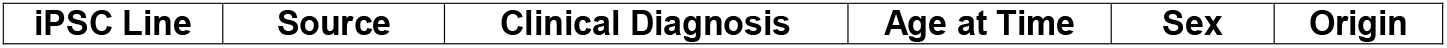
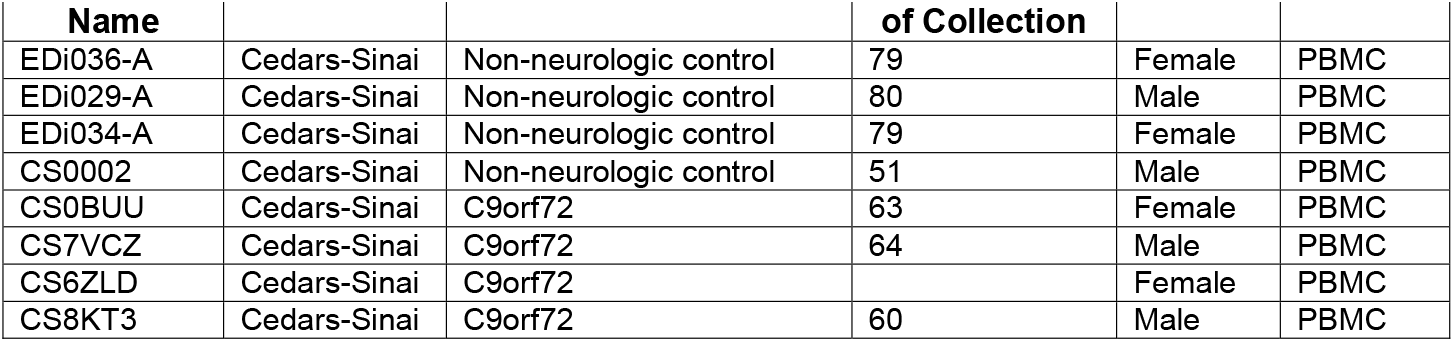
Demographic Information for iPSC Lines.

### ASO Treatment of iPSNs

Scrambled ASO (676630): CCTATAGGACTATCCAGGAA, G_4_C_2_ ASO (619251): CAGGCTGCGGTTGTTTCCCT, and G_2_C_4_ ASO (813214): TCTCATTTCTCTGACCGAAG were generously provided by Ionis Pharmaceuticals. On day 46 of differentiation, ASOs were added to iPSN culture media at a final concentration of 5 μM. On day 49, media was exchanged for fresh stage 3 iPSN culture media and media was subsequently exchanged every 2-3 additional days without ASO until experiments performed at indicated time points.

### DNA Constructs

pcDNA3.1(+) containing 108 sense G_4_C_2_ RNA only repeats was a kind gift from Adrian Isaacs ^31^. To generate an antisense G_2_C_4_ RNA only repeat construct, 108 sense G_4_C_2_ RNA only repeats were digested out of the pcDNA3.1(+) 108 RNA only repeat construct with BamHI and XbaI and inserted in an antisense orientation into empty pcDNA3.1(+) that had been digested with NheI and BamHI. One repeat unit was lost during cloning, resulting in a 107 antisense G_2_C_4_ RNA only repeat construct. Repeat size was confirmed using sequencing with dGTP (Source Bioscience). Sequences are available on request.

### Expression of Repeat RNAs

On day 18 of differentiation, control iPSNs were dissociated with accutase and 5 × 10^6^ iPSNs were transfected with 4 μg G_4_C_2_ and G_2_C_4_ repeat RNA only plasmids in suspension with the Lonza 4D nucleofection system as previously described ^11^. qRT-PCR and immunostaining and confocal imaging were carried out on day 32 as detailed below.

### qRT-PCR

iPSNs were harvested in 1x DPBS with calcium and magnesium and pelleted using a microcentrifuge. 350 μL RLT Buffer was added to iPSN pellets and RNA was isolated using the RNeasy kit (QIAGEN). RNA concentrations were determined using a NanoDrop 1000 spectrophotometer (Thermo Fisher Scientific). For detection of G_4_C_2_ and G_2_C_4_ repeat containing transcripts, 1 μg RNA was used for cDNA synthesis using gene specific primers and the Superscript IV First-Strand cDNA Synthesis System (Thermo Fisher Scientific). For detection of TDP-43 mRNA targets, 1 μg RNA was used for cDNA synthesis using random hexamers and the Superscript IV First-Strand cDNA Synthesis System (Thermo Fisher Scientific). All qRT-qPCR reactions were carried out using SYBR Green Master Mix or TaqMan Gene Expression Master Mix (Thermo Fisher) and an Applied Biosystems QuantStudio 3 (Applied Biosystems). Previously described primer/probe sets (see **Supplemental Table 2** for sequences) ^23^ were used to detect G_4_C_2_ and G_2_C_4_ repeat containing transcripts. Previously described primer sets (see **Supplemental Table 2** for sequences) ^27,28^ were used to detect truncated STMN2 and cryptic exon containing mRNA transcripts. TaqMan Gene Expression Assays (see **Supplemental Table 2** for probe information) were used to detect mRNA targets. GAPDH was used for normalization of gene expression.

**Supplemental Table 2:**
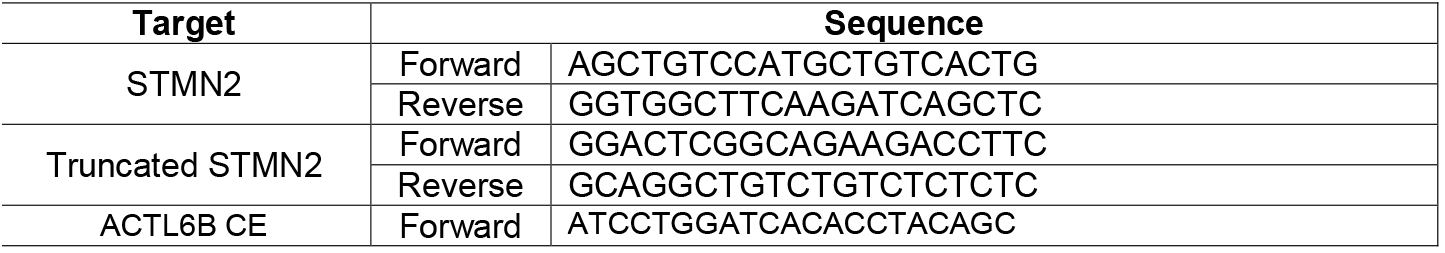
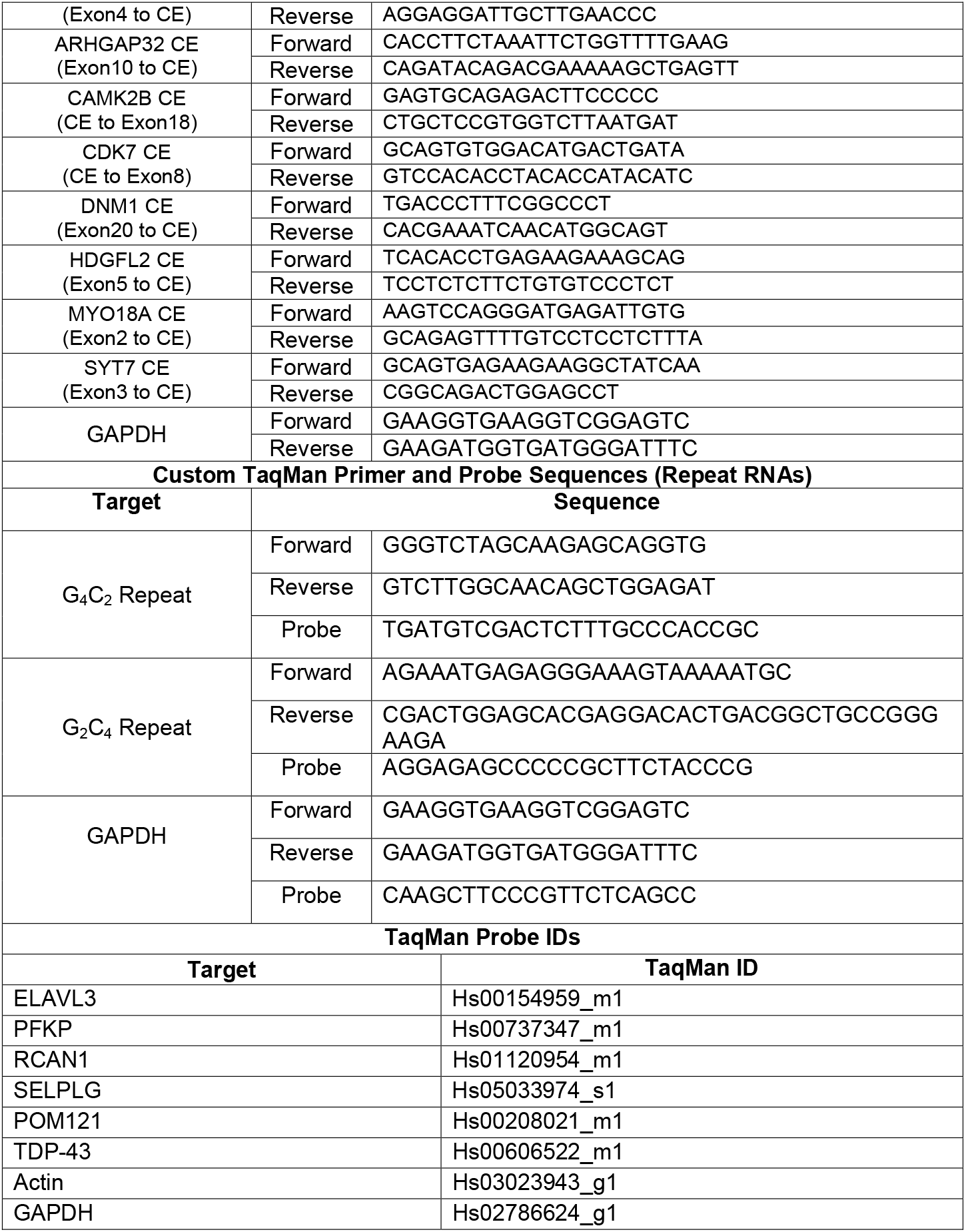
Primer Sequences and TaqMan Probe IDs for qRT-PCR.

### Immunostaining and Confocal Imaging

iPSNs were re-plated using accutase dissociation in Matrigel coated 24 well optical bottom dishes (Cellvis) 3 days prior to fixation. iPSNs were fixed in 4% PFA for 15 minutes, washed 3X 10 minutes with 1X PBS, permeabilized for 15 minutes with 1X PBST containing 0.1% Triton X-100, blocked for 30 minutes in 10% normal goat serum diluted in 1X PBS, and incubated in primary antibody solution (Rabbit Anti-TDP-43 (Proteintech 10782-2-AP), Guinea Pig Anti-Map2 (Synaptic Systems 188004) for 2 hours at room temperature. iPSNs were then washed 3X 10 minutes with 1X PBS, incubated in secondary antibody (Thermo Fisher Scientific; Goat Anti-Rabbit Alexa 488; Goat Anti-Guinea Pig 647) for 1 hour at room temperature, washed 2X 10 minutes in 1X PBS, incubated for 10 minutes with Hoescht diluted 1:1000 in 1X PBS, and washed 2X 10 minutes in 1X PBS. iPSNs were mounted using Prolong Gold Antifade Reagent. iPSNs were imaged using a Zeiss LSM 980 confocal microscope using identical imaging parameters for each well and image. Images presented are maximum intensity projections generated in Zeiss Zen Blue 2.3.

### Immunoprecipitation

On day 32 of differentiation, iPSNs were rinsed in 1X PBS and lysed in 1 mL IP buffer (IP lysis buffer (Thermo Fisher Scientific), 1X protease inhibitor cocktail (Roche), 0.4 U/μL RNasin Plus (Promega)) by scraping with a cell scraper, transferring to an Eppendorf tube, and vortexing for 30-45 seconds. iPSN lysates were centrifuged for 10 minutes at 2500 rpm and 4°C to remove debris. Following centrifugation, the supernatant was transferred to a clean Eppendorf tube. 25 μL of the supernatant was added to 25 μL 2X Laemmli buffer (BioRad) and set aside as protein input and an additional 25 μL supernatant was set aside on ice as RNA input. The remainder of the lysate was split equally between Eppendorf tubes containing 10 μg Rabbit IgG Isotype Control (Thermo Fisher) and Rabbit Anti-TDP-43 (ProteinTech) antibody. Lysates were incubated with antibody with end over end rotation for 2 hours at 4°C. After 2 hours, Magnetic Dynabeads Protein G were washed with IP lysis buffer and 50 μL resuspended bead solution was added to each tube of antibody/lysate solution. Lysate-antibody-bead solutions were incubated with end over end rotation for 2 hours at 4°C. 25 μL of the unbound supernatant was collected and added to 25 μL 2X Laemmli buffer for western blot analysis. The remainder of the unbound supernatant was discarded and immunoprecipitated complexes were washed 3X 5 minutes in 200 μL IP buffer with end over end rotation at 4°C. Bead – IP complexes were resuspended in 200 μL IP buffer. 100 μL of each bead – IP complex sample was set aside for RNA analysis. For the remaining 100 μL, the supernatant was discarded and 50 μL 1X Laemmli buffer was added to bead – IP complexes. All RNA samples (including input) were added to 500 μL RLT Buffer (QIAGEN) and RNA isolation, cDNA synthesis, and qRT-PCR were performed as described above.

### Western Blot

Following immunoprecipitation, samples were heated at 100°C for 5 minutes, and equal volumes of each sample (12.5 μL) were loaded in 4-20% acrylamide gels (BioRad). Gels were run until the dye front reached the bottom. Protein was transferred onto a nitrocellulose membrane using the Trans-Blot Turbo Transfer System (BioRad). Blots were blocked for 30 minutes with 5% nonfat milk in 1X TBST (0.1% Tween-20) and incubated overnight at 4°C with Mouse Anti-TDP-43 (ProteinTech) primary antibody diluted in block. The next day, blots were washed 3X 10 minutes with 1X TBST and probed with Horse Anti-Mouse IgG HRP (Cell Signaling) secondary antibody diluted in block for 1 hour at room temperature. Blots were then washed 3X 10 minutes with 1X TBST and ECL substrate (Thermo Fisher Scientific) was applied for 30 seconds. Chemiluminescent images were acquired with the GE Healthcare ImageQuant LAS 4000 system.

### Statistical Analysis

All image analysis was either completely automated or blinded. Statistical analyses were performed using GraphPad Prism version 9 (GraphPad). For SIM analysis of nuclear POM121 spots, the average of all nuclei evaluated per each iPSC line represents n = 1. The total number of nuclei evaluated per experiment is indicated in the figure legends. Two-way ANOVA with Tukey’s multiple comparison test was used as described in figure legends. * p < 0.05, ** p < 0.01, *** p < 0.001, **** p < 0.0001. Violin plots are used to display the full spread and variability of data within SIM analysis of nuclei. Center dotted line indicates median value. Two additional dotted lines indicate the 25^th^ and 75^th^ percentiles. Bar graphs with individual data points representing each iPSC line are used to display data obtained from all other assays.

## Acknowledgements

We thank the ALS patients and their families for essential contributions to this research. iPSC lines acquired through the Answer ALS program made this research possible. These and other ALS iPSC lines can be obtained from: https://csbiomfg.com/cellcollection. This work was supported by The Robert Packard Center for ALS Research (ANC), NIH NINDS/NIA R00 NS123242 (ANC), Target ALS (ANC, JDR, and PJN), along with funding from NIH-NINDS, NIH-NIA, Department of Defense, ALS Association, Muscular Dystrophy Association, F Prime, and the Chan Zuckerberg Initiative.

## Author Contributions

Conceived and designed the experiments: ANC and JDR. Performed the experiments: ANC, VB, SR, EM. Analyzed the data: ANC. Contributed reagents and materials: ANC, PHN, FR, FB, SM, AMI, and JDR. Wrote the manuscript: ANC and JDR with input from co-authors.

## Figure Legends

**Supplemental Figure 1: TDP-43 cytoplasmic immunoreactivity is observed in G**_**2**_**C**_**4**_ **but not G_4_C_2_ repeat RNA expressing iPSNs**. (**A**) Maximum intensity projections from confocal imaging of TDP-43 in iPSNs 2 weeks following expression of G_4_C_2_ or G_2_C_4_ repeat RNA-only plasmids. Overexpression as indicated on the left, antibody and stain as indicated on top. (**B**) Quantification of the percentage of iPSNs displaying cytoplasmic TDP-43 immunoreactivity. n = 4 control iPSC lines. *** p < 0.001. Scale bar = 10 μm.

**Supplemental Figure 2: G**_**4**_**C**_**2**_ **and G**_**2**_**C**_**4**_ **targeting ASOs significantly reduce pathologic repeat RNA expression in C9orf72 iPSNs**. (**A**) qRT-PCR for G_4_C_2_ repeat RNA in C9orf72 iPSNs 5, 10, 15, and 20 days following a single dose of 5 μM scrambled or repeat targeting ASO. (**B**) qRT-PCR for G_2_C_4_ repeat RNA in C9orf72 iPSNs 5, 10, 15, and 20 days following a single dose of 5 μM scrambled or repeat targeting ASO. GAPDH was used for normalization. N = 4 C9orf72 iPSC lines.

**Supplemental Figure 3: TDP-43 cytoplasmic immunoreactivity is abrogated by G**_**2**_**C**_**4**_ **repeat targeting ASO treatment in C9orf72 iPSNs**. (**A**) Maximum intensity projections from confocal imaging of TDP-43 in iPSNs following 20 days of treatment with G_4_C_2_ or G_2_C_4_ repeat RNA targeting ASOs. ASO treatment as indicated on left, antibody and stain as indicated on top. (**B**) Quantification of the percentage of iPSNs displaying cytoplasmic TDP-43 immunoreactivity. n = 3 control and 3 C9orf72 iPSC lines. ** p < 0.01. Scale bar = 10 μm.

**Supplemental Figure 4: G**_**2**_**C**_**4**_ **repeat RNA is present in TDP-43 complexes in C9orf72 iPSNs**. (**A**) Western blot for TDP-43 following immunoprecipitation. (**B**) qRT-PCR for RNAs following TDP-43 immunoprecipitation in C9orf72 iPSNs. GAPDH was used for normalization. TDP-43 mRNA was used as a positive control. n = 3 C9orf72 iPSC lines. * p < 0.05, **** p < 0.0001.

**Supplemental Figure 5: G_4_C_2_ and G**_**2**_**C**_**4**_ **repeat RNAs are reduced following simultaneous dual ASO treatment**. (**A**) qRT-PCR for G_4_C_2_ repeat RNA in C9orf72 iPSNs 15 days following 5 μM dosing with a combination of scrambled, G_4_C_2_, and G_2_C_4_ repeat RNA targeting ASOs. (**B**) qRT-PCR for G_2_C_4_ repeat RNA in C9orf72 iPSNs 15 days following 5 μM dosing with a combination of scrambled, G_4_C_2_, and G_2_C_4_ repeat RNA targeting ASOs. n = 4 C9orf72 iPSC lines. One-way ANOVA with Tukey’s multiple comparison test was used to calculate statistical significance. **** p < 0.0001.

## References

1 DeJesus-Hernandez, M. et al. Expanded GGGGCC hexanucleotide repeat in noncoding region of C9ORF72 causes chromosome 9p-linked FTD and ALS. Neuron 72, 245–256, doi:10.1016/j.neuron.2011.09.011 (2011).

2 Renton, A. E. et al. A hexanucleotide repeat expansion in C9ORF72 is the cause of chromosome 9p21-linked ALS-FTD. Neuron 72, 257–268, doi:10.1016/j.neuron.2011.09.010 (2011).

3 Gitler, A. D. & Tsuiji, H. There has been an awakening: Emerging mechanisms of C9orf72 mutations in FTD/ALS. Brain research 1647, 19–29, doi:10.1016/j.brainres.2016.04.004 (2016).

4 Gupta, R. et al. The Proline/Arginine Dipeptide from Hexanucleotide Repeat Expanded C9ORF72 Inhibits the Proteasome. eNeuro 4, pdoi:10.1523/eneuro.0249-16.2017 (2017).

5 Lin, Y. et al. Toxic PR Poly-Dipeptides Encoded by the C9orf72 Repeat Expansion Target LC Domain Polymers. Cell 167, 789–802.e712, doi:10.1016/j.cell.2016.10.003 (2016).

6 Wen, X. et al. Antisense proline-arginine RAN dipeptides linked to C9ORF72-ALS/FTD form toxic nuclear aggregates that initiate in vitro and in vivo neuronal death. Neuron 84, 1213–1225, doi:10.1016/j.neuron.2014.12.010 (2014).

7 Zhang, Y. J. et al. Heterochromatin anomalies and double-stranded RNA accumulation underlie C9orf72 poly(PR) toxicity. Science (New York, N.Y.) 363, pdoi:10.1126/science.aav2606 (2019).

8 Freibaum, B. D. & Taylor, J. P. The Role of Dipeptide Repeats in C9ORF72-Related ALS-FTD. Frontiers in molecular neuroscience 10, 35, doi:10.3389/fnmol.2017.00035 (2017).

9 Conlon, E. G. et al. The C9ORF72 GGGGCC expansion forms RNA G-quadruplex inclusions and sequesters hnRNP H to disrupt splicing in ALS brains. eLife 5, pdoi:10.7554/eLife.17820 (2016).

10 Zhang, K. et al. The C9orf72 repeat expansion disrupts nucleocytoplasmic transport. Nature 525, 56–61, doi:10.1038/nature14973 (2015).

11 Coyne, A. N. et al. G(4)C(2) Repeat RNA Initiates a POM121-Mediated Reduction in Specific Nucleoporins in C9orf72 ALS/FTD. Neuron, doi:10.1016/j.neuron.2020.06.027 (2020).

12 Vatsavayai, S. C., Nana, A. L., Yokoyama, J. S. & Seeley, W. W. C9orf72-FTD/ALS pathogenesis: evidence from human neuropathological studies. Acta neuropathologica 137, 1–26, doi:10.1007/s00401-018-1921-0 (2019).

13 Vatsavayai, S. C. et al. Timing and significance of pathological features in C9orf72 expansion-associated frontotemporal dementia. Brain : a journal of neurology 139, 3202–3216, doi:10.1093/brain/aww250 (2016).

14 Cooper-Knock, J. et al. Antisense RNA foci in the motor neurons of C9ORF72-ALS patients are associated with TDP-43 proteinopathy. Acta neuropathologica 130, 63–75, doi:10.1007/s00401-015-1429-9 (2015).

15 Davidson, Y. S. et al. Brain distribution of dipeptide repeat proteins in frontotemporal lobar degeneration and motor neurone disease associated with expansions in C9ORF72. Acta neuropathologica communications 2, 70, doi:10.1186/2051-5960-2-70 (2014).

16 DeJesus-Hernandez, M. et al. In-depth clinico-pathological examination of RNA foci in a large cohort of C9ORF72 expansion carriers. Acta neuropathologica 134, 255–269, doi:10.1007/s00401-017-1725-7 (2017).

17 Gendron, T. F. et al. Antisense transcripts of the expanded C9ORF72 hexanucleotide repeat form nuclear RNA foci and undergo repeat-associated non-ATG translation in c9FTD/ALS. Acta neuropathologica 126, 829–844, doi:10.1007/s00401-013-1192-8 (2013).

18 Lee, Y. B. et al. Hexanucleotide repeats in ALS/FTD form length-dependent RNA foci, sequester RNA binding proteins, and are neurotoxic. Cell reports 5, 1178–1186, doi:10.1016/j.celrep.2013.10.049 (2013).

19 Mackenzie, I. R., Frick, P. & Neumann, M. The neuropathology associated with repeat expansions in the C9ORF72 gene. Acta neuropathologica 127, 347–357, doi:10.1007/s00401-013-1232-4 (2014).

20 Mizielinska, S. et al. C9orf72 frontotemporal lobar degeneration is characterised by frequent neuronal sense and antisense RNA foci. Acta neuropathologica 126, 845–857, doi:10.1007/s00401-013-1200-z (2013).

21 Aladesuyi Arogundade, O. et al. Antisense RNA foci are associated with nucleoli and TDP-43 mislocalization in C9orf72-ALS/FTD: a quantitative study. Acta neuropathologica 137, 527–530, doi:10.1007/s00401-018-01955-0 (2019).

22 Donnelly, C. J. et al. RNA toxicity from the ALS/FTD C9ORF72 expansion is mitigated by antisense intervention. Neuron 80, 415–428, doi:10.1016/j.neuron.2013.10.015 (2013).

23 Lagier-Tourenne, C. et al. Targeted degradation of sense and antisense C9orf72 RNA foci as therapy for ALS and frontotemporal degeneration. Proceedings of the National Academy of Sciences of the United States of America 110, E4530–4539, doi:10.1073/pnas.1318835110 (2013).

24 Gendron, T. F. et al. Poly(GP) proteins are a useful pharmacodynamic marker for C9ORF72-associated amyotrophic lateral sclerosis. Science translational medicine 9, pdoi:10.1126/scitranslmed.aai7866 (2017).

25 Brown, A. L. et al. TDP-43 loss and ALS-risk SNPs drive mis-splicing and depletion of UNC13A. Nature 603, 131–137, doi:10.1038/s41586-022-04436-3 (2022).

26 Klim, J. R. et al. ALS-implicated protein TDP-43 sustains levels of STMN2, a mediator of motor neuron growth and repair. Nature neuroscience 22, 167–179, doi:10.1038/s41593-018-0300-4 (2019).

27 Melamed, Z. et al. Premature polyadenylation-mediated loss of stathmin-2 is a hallmark of TDP-43-dependent neurodegeneration. Nature neuroscience 22, 180–190, doi:10.1038/s41593-018-0293-z (2019).

28 Seddighi, S. et al. Mis-spliced transcripts generate <em>de novo</em> proteins in TDP-43-related ALS/FTD. bioRxiv, 2023.2001.2023.525149, doi:10.1101/2023.01.23.525149 (2023).

29 Ling, J. P., Pletnikova, O., Troncoso, J. C. & Wong, P. C. TDP-43 repression of nonconserved cryptic exons is compromised in ALS-FTD. Science (New York, N.Y.) 349, 650–655, doi:10.1126/science.aab0983 (2015).

30 Coyne, A. N. et al. Nuclear accumulation of CHMP7 initiates nuclear pore complex injury and subsequent TDP-43 dysfunction in sporadic and familial ALS. Science translational medicine 13, doi:10.1126/scitranslmed.abe1923 (2021).

31 Mizielinska, S. et al. C9orf72 repeat expansions cause neurodegeneration in Drosophila through arginine-rich proteins. Science (New York, N.Y.) 345, 1192–1194, doi:10.1126/science.1256800 (2014).

32 Moens, T. G. et al. Sense and antisense RNA are not toxic in Drosophila models of C9orf72-associated ALS/FTD. Acta neuropathologica 135, 445–457, doi:10.1007/s00401-017-1798-3 (2018).

33 Prudencio, M. et al. Misregulation of human sortilin splicing leads to the generation of a nonfunctional progranulin receptor. Proceedings of the National Academy of Sciences of the United States of America 109, 21510–21515, doi:10.1073/pnas.1211577110 (2012).

34 Liu, Y. et al. C9orf72 BAC Mouse Model with Motor Deficits and Neurodegenerative Features of ALS/FTD. Neuron 90, 521–534, doi:10.1016/j.neuron.2016.04.005 (2016).

35 Chew, J. et al. Aberrant deposition of stress granule-resident proteins linked to C9orf72-associated TDP-43 proteinopathy. Molecular neurodegeneration 14, 9, doi:10.1186/s13024-019-0310-z (2019).

36 Baxi, E. G. et al. Answer ALS, a large-scale resource for sporadic and familial ALS combining clinical and multi-omics data from induced pluripotent cell lines. Nature neuroscience 25, 226–237, doi:10.1038/s41593-021-01006-0 (2022).

