## Supplementary figures and images for "G_2_C_4_ targeting antisense oligonucleotides potently mitigate TDP-43 dysfunction in C9orf72 ALS/FTD human neurons"

### Supplemental Figure 1

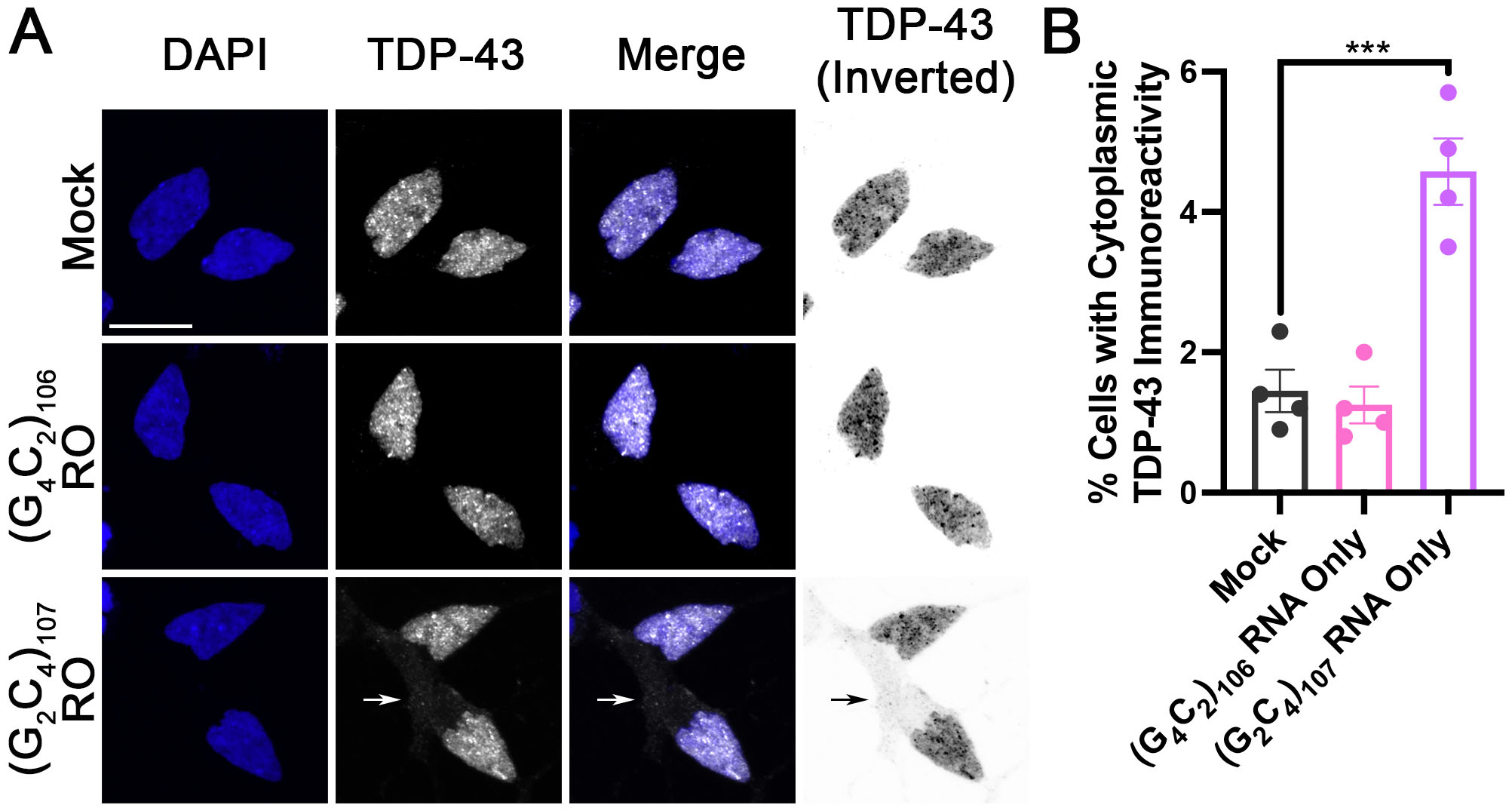

### Supplemental Figure 2

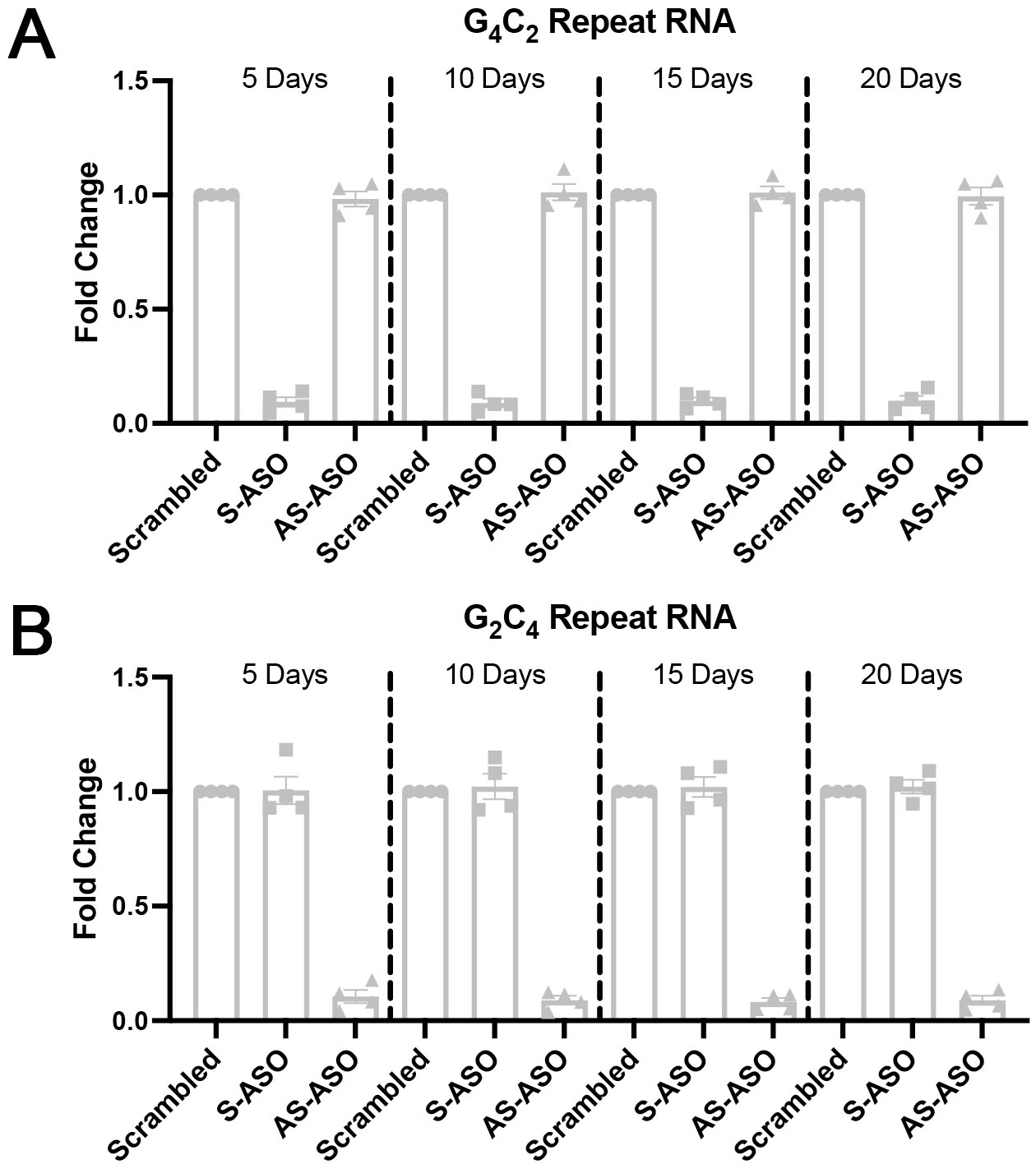

### Supplemental Figure 3

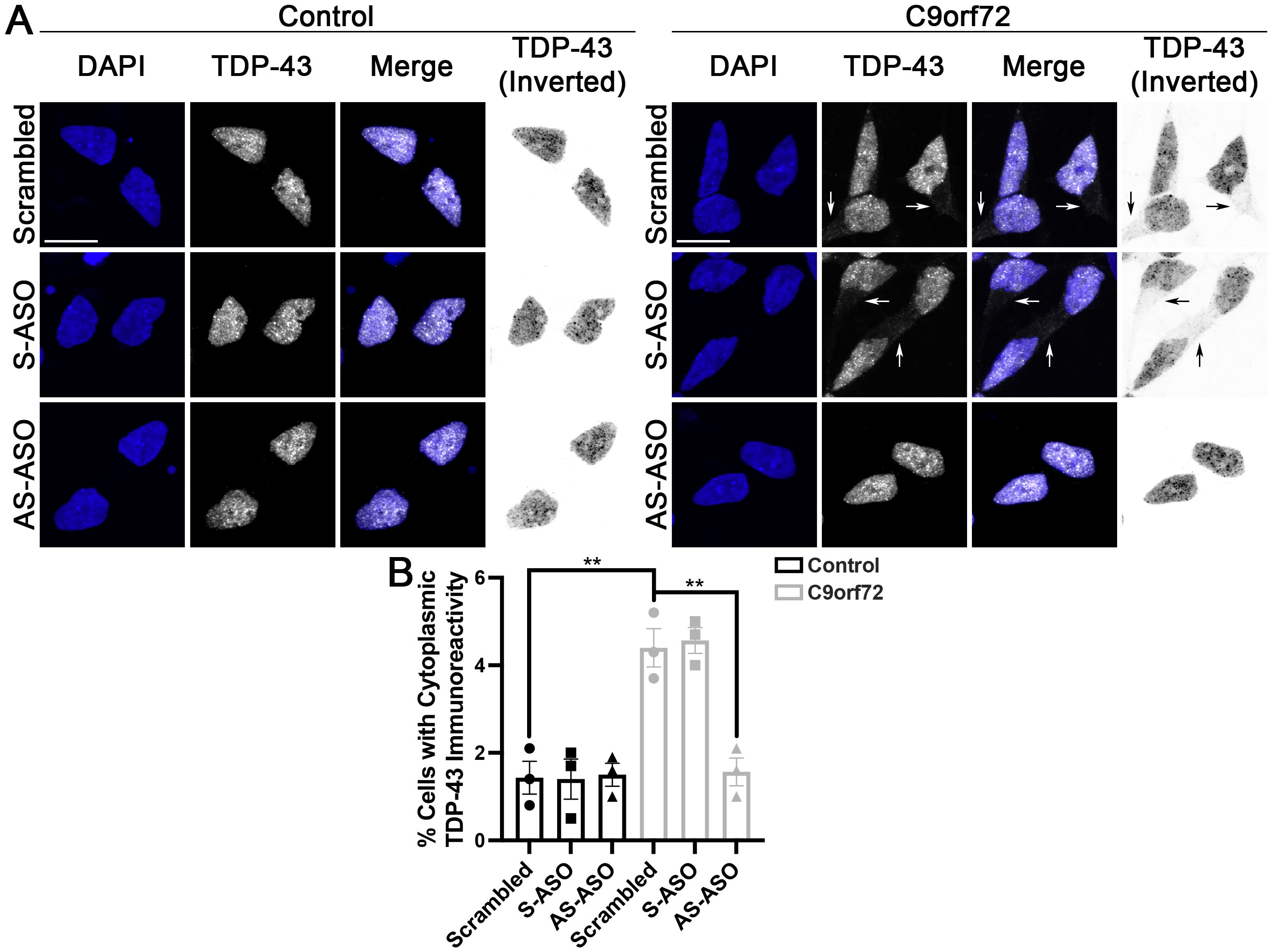

### Supplemental Figure 4

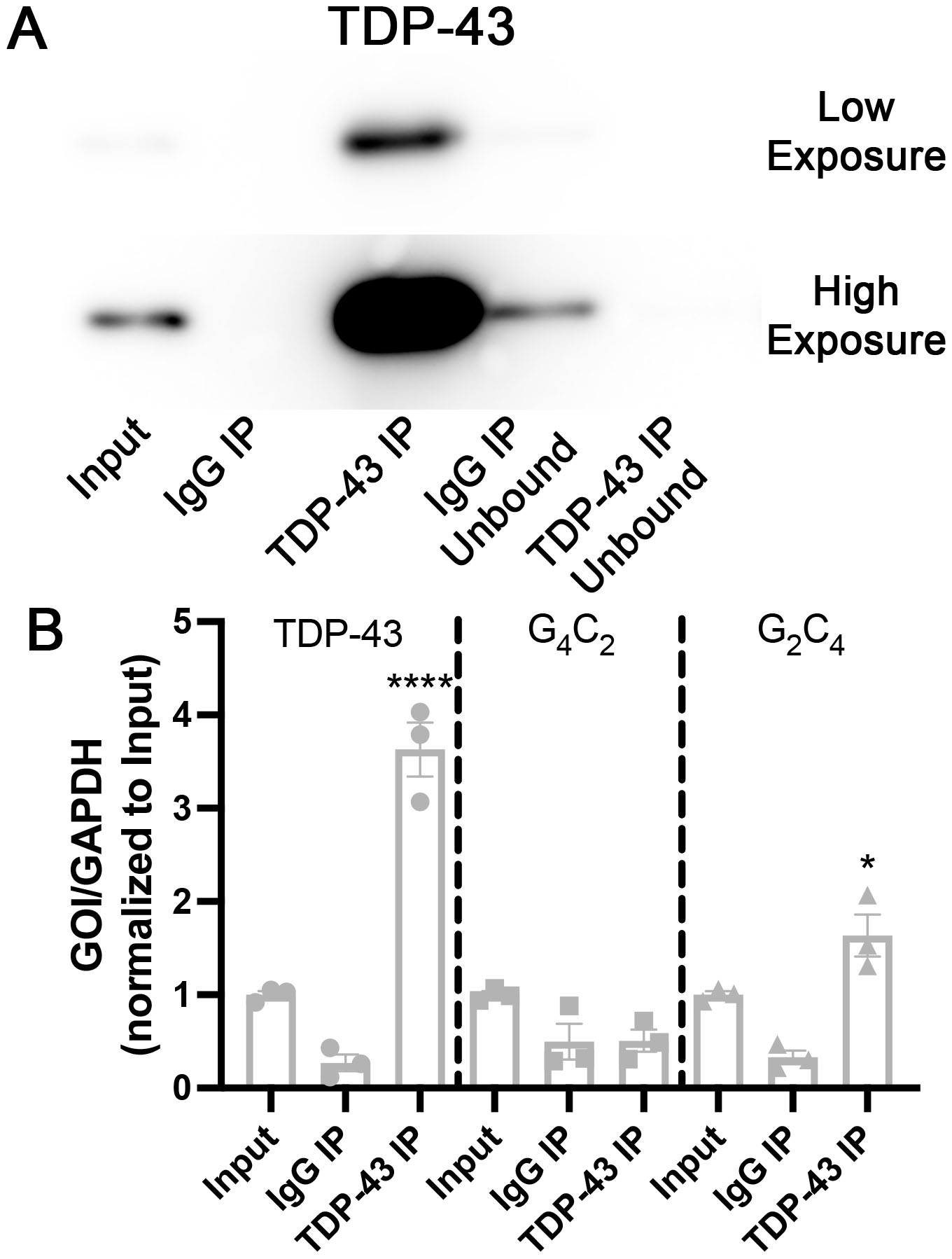

### Supplemental Figure 5

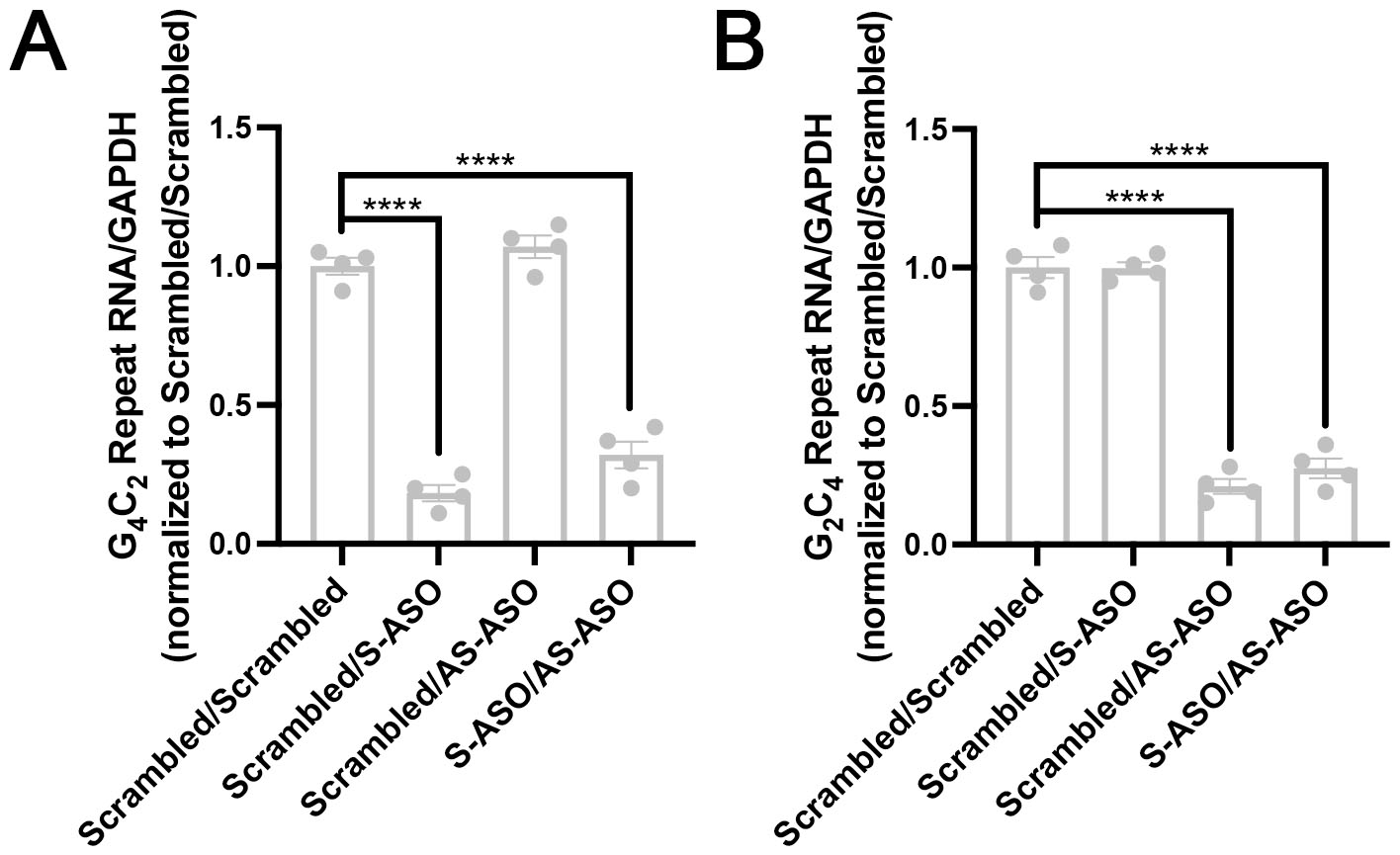
